# Single-residue variation in the nucleosome core reveals a regulatory hub for phenotypic innovation

**DOI:** 10.64898/2026.09.11.750863

**Authors:** Zachary H. Harvey, Benjamin Gundinger, Jian Yi Kok, Kathryn M. Stevens, Tobias Warnecke, Frédéric Berger

**Affiliations:** Gregor Mendel Institute, Austrian Academy of Sciences, Vienna, Austria; MRC Laboratory of Medical Sciences, London, United Kingdom; Institute of Clinical Sciences, Imperial College London, London, United Kingdom; Trinity College, Oxford, United Kingdom; Department of Biochemistry, University of Oxford, Oxford, United Kingdom

## Abstract

Nucleosomes organize genomes and regulate DNA access, yet accumulating evidence suggests that their constituent histones may have functions beyond canonical chromatin regulation, but the breadth of such regulatory diversity remains unclear. Here, we used the six-residue loop 2 (L2) of H2A and H2A.Z to map, at single-residue resolution, how nucleosome-core variation reshapes cellular function. Genome-scale interaction mapping identified hundreds of regulatory connections spanning chromatin, as well as actin organization, endocytosis, and membrane trafficking. Interactions were residue-specific and differed between H2A and H2A.Z, revealing a regulatory landscape encoded by single residues. Transcriptome profiling showed limited expression changes and little overlap between differentially expressed genes and regulatory partners, indicating that non-chromatin connections are not readily explained by altered transcription. L2 substitutions also preferentially conferred benefits under cell wall and membrane stress. Thus, the nucleosome is linked to cellular-periphery functions beyond classical chromatin regulation, identifying histone variation as a source of phenotypic innovation.

## INTRODUCTION

Nucleosomes are ancient complexes of histone proteins and DNA^1,2^ thought to have evolved to solve the challenge of packaging eukaryotic genomes.^3,4^ This fundamental structural role has been linked to essential genome regulation functions, including coordinating gene expression during environmental^5^ and developmental transitions.^6^ However, this view is incomplete. Accumulating evidence indicates that nucleosomes and their histone constituents have diverse functions beyond genome packaging.^7^ These include antimicrobial activity following histone release by neutrophils,^8^ roles in metabolism through the sequestration of methyl groups,^9^ and enzymatic reduction of copper to support cellular homeostasis.^10^ These examples highlight a broader mechanistic repertoire for these highly abundant complexes in addition to chromatin structure, but the scope of such functional diversity remains unclear.

Small changes to the nucleosome’s histones observed across eukaryotic diversity and in human disease suggest a much broader role for the nucleosome. Although nucleosome structure has remained nearly invariant throughout eukaryotic evolution, histones have diversified both between species^11–13^ and within them.^13,14^ This diversification can have substantial biological consequences: single-residue differences among histone gene copies can produce large functional differences,^15,16^ and recurrent mutations in histones in human cancers, collectively termed oncohistones, occur at rates similar to other oncogenic drivers and can drive pathology.^11,12,17–21^ Yet, even oncohistones predicted to disrupt genome packaging or transcription often have limited effects on either process,^11^ indicating that the mechanistic basis of functionally important histone sequence variation remains unresolved.

The histone H2A family provides a powerful model for addressing whether functional variation might reveal additional regulatory scope of the nucleosome. Diverse variants of H2A have evolved,^22^ including H2A.Z that diverged from H2A at the dawn of eukaryotes^1,13^ and lineage-specific variants of H2A such as macroH2A and H2A.W.^23,24^ Although these histone variants have discrete and conserved roles, small sequence changes within each variant lineage still impact their function. For example, despite being ∼90% sequence identical across all eukaryotes,^13^ H2A.Z can be either a transcriptional activator in some species,^25^ or a repressor in others;^26^ a functional difference distinguished by single-residue changes to the loop 2 (L2) that links the second two alpha helices of its core domain.^13^ Further, both H2A.Z and H2A have diverse functional roles besides transcription in processes including chromosome segregation,^27^ mRNA splicing^28^ and genome maintenance,^13,29–32^ but their mechanistic basis remains largely unknown.

Using the H2A/H2A.Z L2 region as a molecular entry point, here we show that the nucleosome’s role extends far beyond regulating the genome. Using high-resolution genetic mapping, we demonstrate that the L2 forms a dense network of interactions with processes as diverse as transcription, translation, metabolism, endocytosis, and the actin cytoskeleton. Such regulatory diversity is encoded with single-residue precision, with neighboring residues in both H2A.Z and H2A having divergent regulatory impact. Further, we show that the L2 regulates functional categories across transcription, actin, endocytosis, and membrane trafficking, and that non-transcriptional functions are not an indirect effect of transcriptional changes. Finally, we demonstrate that diverse regulatory pathways are anchored to specific L2 residues. Together, our results expand the regulatory repertoire of the nucleosome, suggesting that some of its oldest and most conserved functions lie outside of canonical gene regulation and chromatin architecture.

## RESULTS

### The nucleosome core L2 encodes phenotypic diversity

To understand the nucleosome’s regulatory function, we focused on the H2A/H2A.Z loop 2 (L2), which is the principal determinant of H2A and H2A.Z’s functional divergence.^13,32^ Whereas the L2 does not directly impact nucleosome stability,^13^ it is among the most highly solvent-accessible region on the side face of the nucleosome (Fig. 1A blue trace, B). We previously reported that this region directly contacts the transcriptional regulator Spt6.^13^ However, this interaction was primarily related to the molecular function of a single residue of H2A.Z and did not clarify H2A.Z’s broader genomic functions.^23,27,33^ Further, the L2 has among the highest sequence diversity across H2A/H2A.Z in both natural and disease-associated sequence variants (Fig. 1A black trace, B; Fig. S1A), suggesting that these 6 residues could have regulatory complexity beyond its reported role in transcription.^13^

**Figure 1.**
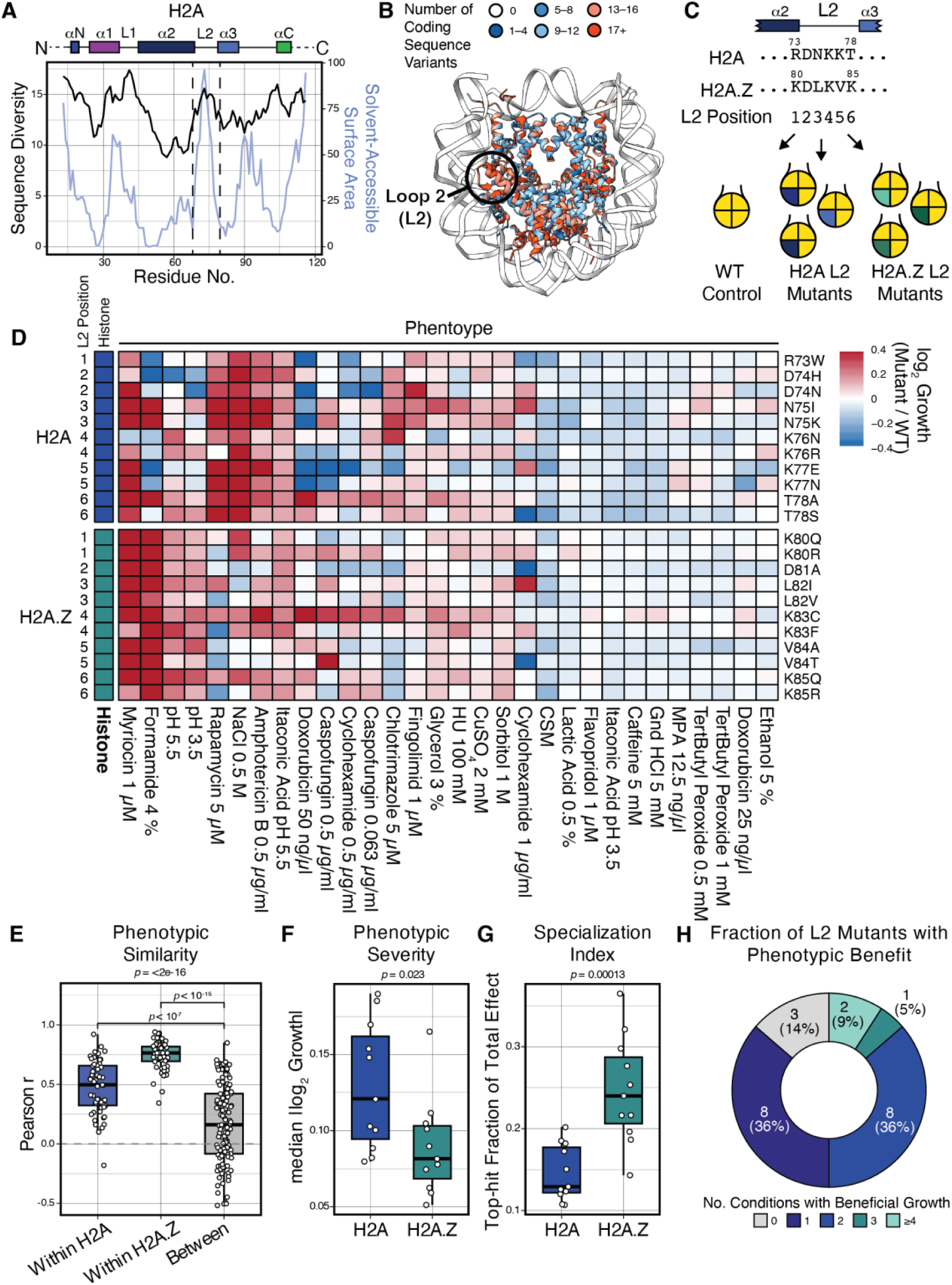
The nucleosome core L2 encodes phenotypic diversity. A, Sequence diversity of H2A is calculated from a database of H2A sequences from eukaryotic diversity^13^ and human cancers.^12^ Sequence diversity is defined as the number of unique alternative amino acids present in all sequences in our database, and is presented by a 3-amino acid rolling average. Solvent accessibility per residue was calculated with ChimeraX from PDB 1ID3. **B,** Sequence diversity (A) projected onto the yeast nucleosome structure (PDB 1ID3). **C,** Scheme illustrating the library of L2 mutants used in this study. **D,** Phenotypic diversity of L2 mutants. L2 position and histone backbone for each mutant is added, and growth is expressed as log2 area under the curve for the mutant relative to the WT in a given stress condition. Abbreviated conditions: HU, hydroxyurea; Gnd HCl, guanidinium hydochloride; MPA, mycophenolic acid. **E,** Correlation analysis of L2 mutants either within or between histones. Statistical testing was done with a Wilcoxon rank test, except for the three-way comparison, which is a Kruskal-Wallis test. **F,** Comparison of H2A and H2A.Z L2 mutant phenotypic severity, defined as the median of the absolute growth effect across all assayed conditions. **G,** Comparison of H2A and H2A.Z specialization index. The specialization index is the fraction of the total phenotypic severity per L2 position that is explained by the strongest phenotype within a condition. For both (F) and (G) statistical significance (*p*) was calculated with a Wilcoxon rank test. **H,** Fraction of L2 mutants with beneficial phenotypes, defined as >50% growth advantage relative to WT.

To test the potential breadth of the L2’s function, we systematically mutated each of its 6 residues across both H2A and H2A.Z to 1–2 biochemically distinct substitutions selected either from eukaryotic diversity^13^ or cancer-associated mutations,^12^ in total generating 22 different single residue substitutions (Fig. 1C). To assess their impact on the nucleosome’s function, we first assessed their effect on the cell’s response to a broad spectrum of stress conditions spanning both general growth and environmental conditions, as well as clinically relevant pharmacological perturbations. These included antifungal drugs (caspofungin, clotrimazole, amphotericin B), osmotic stressors (NaCl and sorbitol), DNA replication and genotoxic stressors (hydroxyurea and doxorubicin), translational and proteotoxic stressors (cycloheximide and guanidinium hydrochloride), inhibitors of transcription (flavopiridol, mycophenolic acid (MPA)) and of membrane dynamics (fingolimod, myriocin), as well as metabolic (glycerol, rapamycin) and oxidative (tert-butyl hydroperoxide) challenges, and general stressors (ethanol, formamide, copper sulfate, caffeine, lactic acid, itaconic acid, and acidic pH conditions).

Across the stress panel, H2A and H2A.Z mutants exhibited diverse fitness profiles (Fig. 1D). Consistent with their evolutionary diversification, H2A and H2A.Z’s phenotypic profiles differed markedly from one another; a given H2A mutant was phenotypically more similar to another H2A mutant than an H2A.Z mutant, and vice versa (*p* < 2.2 x 10^-16^, Kruskal-Wallis; Fig. 1D–E, S1A). Finally, H2A.Z’s phenotypes were overall relatively milder than H2A (Fig. 1D, F). However, whereas H2A tended to have global phenotypes whereby a given L2 mutant affected growth across multiple conditions, those for H2A.Z tended to be more specialized: specific L2 mutants had strong phenotypic effects concentrated in fewer conditions (Fig. 1G; S1B–C). Finally, whereas ∼82% of loss-of-function alleles were previously shown to be phenotypically detrimental in *Saccharomyces cerevisiae*,^34^ we were surprised that 86% of all tested L2 mutants were phenotypically beneficial in at least one condition (growth >50% greater than WT; Fig. 1H). Together, these data establish that the L2 is an exquisitely sensitive determinant of phenotypic potential, that H2A and H2A.Z markedly diverge in their function, and that H2A.Z tends to be more specialized.

### The nucleosome core L2 is a regulatory hub with single-residue precision

We next asked what the regulatory basis for the L2’s phenotypic diversity might be. To do this, we mapped the genetic interactions of each of L2 single residue substitution mutants, and WT controls, using a variant of the synthetic genetic array (SGA) approach^35^ (pE-MAP^36^). Each of the L2 single residue substitutions, and WT controls, were crossed to both the yeast knock-out^37^ and knock-down (DAmP^38^) collections to construct a systematic library of double mutants combining one L2 substitution with one gene disruption (Fig. 2A). This yielded 134,527 unique double mutant pairs, allowing us to assess the genetic dependency of each L2 mutant to ∼90% of all ORFs in the *S. cerevisiae* genome. Using this library of unique double mutants, we assayed growth across four diverse conditions (rich medium, as well as salt, heat, and genotoxic stress). Conditions were chosen based on their large-magnitude phenotypic effect, focusing on general stress conditions with distinct profiles across different L2 mutants (Fig. 1D). In total, we measured ∼1 million datapoints, allowing us to calculate genetic interaction scores (GIs) for all unique double mutant pairs. A GI was defined as the quantitative deviation in growth of a given double mutant relative to the two individual mutations alone. In this analysis, a negative GI corresponds to synthetic sick/lethal double mutants, where a positive GI corresponds to synthetic viable/rescue double mutants. GI values were calculated by applying a linear mixed effects model to account for both biological and technical variation (Fig. S2A). Using a stringent cutoff (|GI| > 0.3 and |Z| > 2.5), we identified 289 total GIs across both H2A and H2A.Z (Data S1), which together represent a comprehensive regulatory network of the L2.

**Figure 2.**
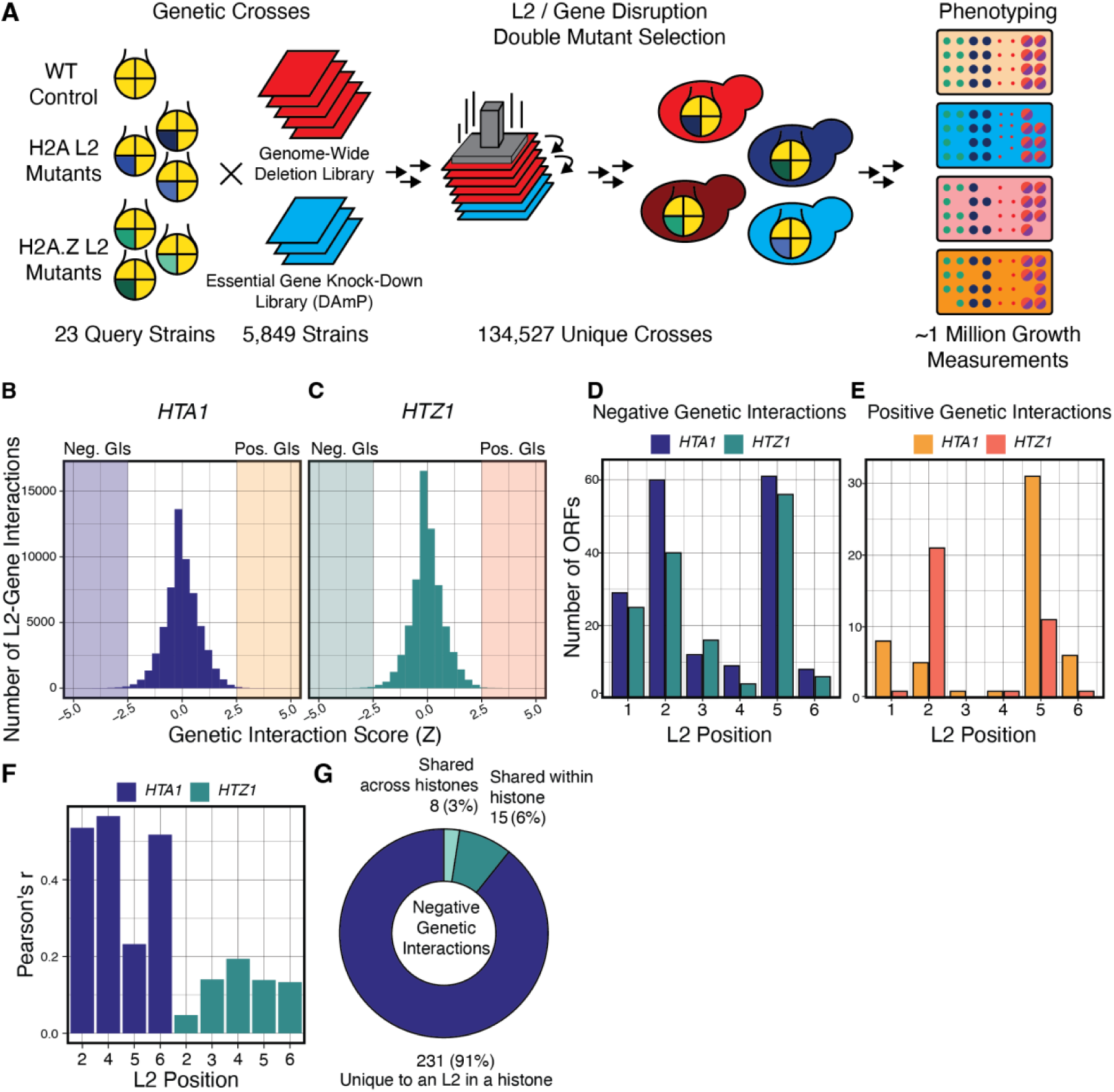
The nucleosome core L2 is regulatory hub at single-residue precision. A, Experimental scheme for mapping the comprehensive genetic interactions of each L2 position of H2A and H2A.Z. **B–C,** Distribution of genetic interaction (GI) scores for both H2A (*HTA1*; B) and H2A.Z (*HTZ1*; C) across all 21 assayed L2 mutants. **D–E,** Number of GIs per H2A and H2A.Z position for both negative (D) and positive (E) interactions. **F,** Correlation analysis (Pearson’s r) of GIs by L2 position and histone. For each position, all GI scores for distinct amino acid substitutions at that position were compared, except for those positions that had only one mutation, which were excluded. **G,** L2 mutant specificity of negative genetic interactions. The pie chart represents the fraction of negative GIs that were unique to a specific L2 position in a given histone versus those shared either within or between histones.

We first compared our identified GIs to the systematic genetic interaction map between H2A and H2A.Z disruptions in *S. cerevisiae*.^39–41^ Consistent with previous observations of the genetic characteristics of single-residue substitutions relative to entire gene deletions,^36^ we identified minimal overlap between our L2 mutant GIs and those of either *hta1*Δ or *htz1*Δ (Fig. S2B). This indicates that histones’ gene deletion mutations are likely dominated by compensatory network effects, whereas single-residue substitutions do not phenocopy loss-of-function but rather disrupt specific interactions. GIs were not uniformly distributed across the L2, but were concentrated at L2 positions 1, 2, and 5 (Fig. 2D–E). Whereas distinct substitutions at a given L2 position in H2A had broadly similar GIs, H2A.Z GIs depended not only on L2 position, but also amino acid identity (Fig. 2F; S2C). This is consistent with H2A.Z’s phenotypic effects in specific growth conditions (Fig. 1G; S1B–C), and further reflective of H2A.Z’s specialization for specific regulatory contexts.^42^ Finally, the vast majority of GIs (91%) were specific to a given L2 position and histone, with only 2% shared between them (Fig. 2G; S2D). Together, these data indicate that the 6-residue L2 region is a dense regulatory region with specific interactions defined by single residues.

### The nucleosome core encodes regulatory diversity for nuclear and non-nuclear functions

We next asked how the L2’s genetic complexity was arranged into regulatory networks. To do this, we used STRING^43^ annotations to construct a functional interaction network from all L2 negative (Fig. 3A) and positive (Fig. 3B) GIs across both H2A and H2A.Z. These networks were highly coherent (*p* < 0.001, permutation test). As expected, a substantial proportion (45%/53% for negative and positive GIs, respectively) of the L2 GI networks’ nodes were linked to chromatin (Fig. 3C–D). However, whereas the GIs for complete loss-of-function alleles (i.e. *hta1*Δ and *htz1*Δ^39^) were completely dominated by a general enrichment for chromatin-related genes (*p*_adj_ < 10^-16^, Fisher’s exact test; Fig. S3A–B), GIs for L2 mutants were not (Fig. S3C–D). Rather, consistent with our prior observations,^13^ L2 genetic networks were specifically enriched for transcription-related genes (*p*_adj_ < 0.05, Fisher’s exact test; Fig. S3E). This comprised central components of RNA polymerase II (RNAPII) gene transcription, including both its subunit B12.5 (*RPB11*) and transcription factor II F (TFIIF/*TFG1*). This suggests that the L2 point mutants are separation-of-function mutants specifically impacting transcription without grossly perturbing general chromatin structure.^13^

**Figure 3.**
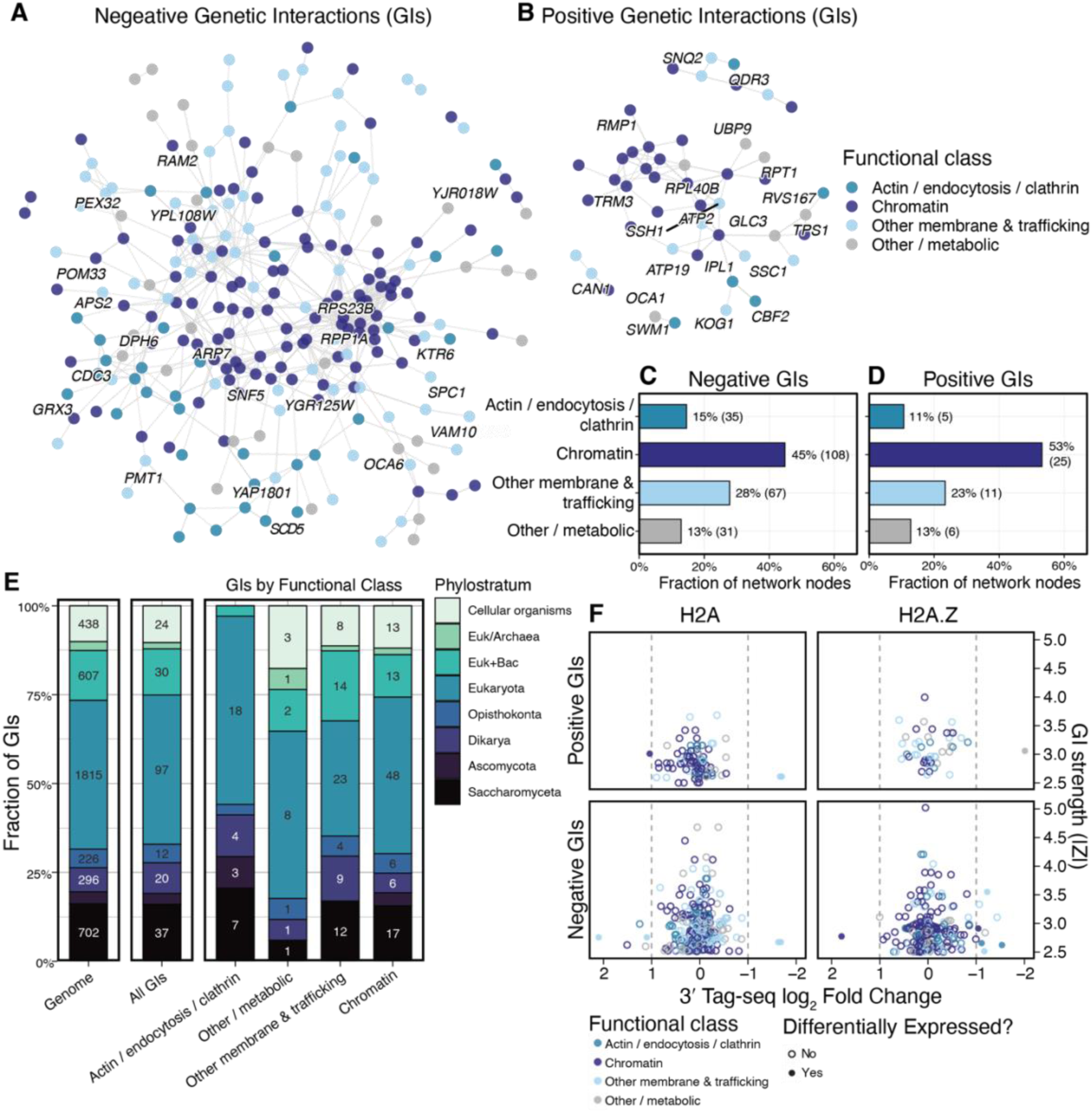
The nucleosome core encodes regulatory diversity for nuclear and non-nuclear functions. A–B, STRING network for negative (A) or positive (B) genetic interactions. Each node is a GI for an L2 mutant, and is colored based on GO annotation across four assayed categories (actin, endocytosis, and clathrin; Transcription and chromatin; other membrane-related functions; other & metabolic; see methods). Orphan nodes were pruned, and for each of the four annotation categories, the top GIs within are labeled. **C–D,** Breakdown of the number of GIs per functional category from STRING networks. **E,** GIs by gene age, as calculated by Liebeskind, et al.^46^ For reference, the age distribution of all genes in the *Saccharomyces cerevisiae* genome is provided, with the number of genes in each category given within each box. **F,** Comparison of L2 GIs with their transcriptional impact. For each high-confidence GI, the log_2_ fold change in expression from 3’ Tag-seq experiments in matched L2 mutants is plotted. Points are colored by functional category, and significantly differentially expressed (*p*_adj_ < 0.05 and |log_2_ fold change| > 1) genes are filled.

We were surprised that nearly half of GIs (43% total) were linked to non-chromatin functions, including actin and endocytosis (15%), as well as general membrane dynamics and trafficking (28%; Fig. 3A–D; Data S1). Whereas membrane and trafficking-related interactions were not significantly enriched above the genome-wide average, GIs for actin, endocytosis, and clathrin-related proteins were (*p*_adj_ < 0.01, Fisher’s exact test; Fig. S3A,F). Importantly, actin and endocytosis enrichments were not driven by actin-related proteins known to be involved in diverse chromatin remodeling complexes (Data S1).^44^ Reinforcing this functional enrichment, when we examined beneficial phenotypes (>50% growth advantage relative to WT), L2 mutants were specifically enriched for stresses that perturb cell-wall or membrane homeostasis (overall OR= 3.32, *p* < 0.001, Fisher’s exact test), with a slightly higher enrichment for H2A.Z than for H2A (OR = 4.0, *p* < 0.01, Fisher’s exact test; Fig. S4A). Further, consistent with a specific impact on non-chromatin functions, phenotypic benefit was strongest in the sphingolipid inhibitor myriocin (Fig. S4B, 1D), which specifically perturbs membranes, and somewhat weaker for antifungal drugs that only indirectly impact membrane dynamics (Fig. S4C–E, 1D). Such stresses are directly linked to the molecular function of actin and endocytosis, which together are important remodelers of the plasma membrane and cell wall.^45^ Whereas these drug-response phenotypes are specific to fungi, we next asked whether the genetic interactors driving these phenotypes were too. To do this, we examined the phylogenetic age^46^ of both genetic interactors overall or broken down by category (Fig. 3E). Like GIs across all categories and the *S. cerevisiae* genome, the majority of actin, endocytosis, and clathrin related GIs were highly conserved across all eukaryotes (Fig. 3E). This suggests that the L2’s regulatory network is conserved. Together, these data identify a previously unappreciated functional connection of histones to the cellular periphery.

We wondered whether functional links across actin and endocytosis, or other categories like membrane trafficking or metabolism could be explained by misexpression caused by the L2’s impact on transcription.^13,32^ If this were the case, then L2 mutants should have significant differential expression of genes linked to these non-canonical functions, including those genes that emerged as GIs. To test this, we profiled the transcriptomes of L2 mutants, comparing their differentially expressed genes relative to matched WT controls (Fig. S4A). Overall, transcriptomic changes were limited among L2 mutants (Fig. S4B). Many L2 substitutions produced few or no significantly differentially expressed genes relative to matched WT controls (Fig. S4A–B), and those few differentially expressed genes had no common GO term enrichments indicative of a shared functional role. To assess whether, despite their low numbers, differentially expressed genes could still explain observed GIs, we examined whether GIs were differentially expressed. We compared negative and positive GIs to differentially expressed genes for each H2A and H2A.Z L2 mutant. Almost none of the L2 GIs were differentially expressed (Fig. 3F; S4C). To account for the possibility that, although the GIs themselves are not differentially expressed, those genes upstream or downstream of them in their genetic pathways were, we checked whether genes associated with the cell wall phenotypes we observed were differentially expressed. We used curated phenotypic data from the *Saccharomyces* genome database (SGD)^34^ to assemble a list of genes known to either increase or decrease growth in response to the assayed cell wall and membrane stressors (i.e. the antifungal drugs amphotericin B, caspofungin, clotrimazole, and the sphingolipid inhibitors fingolimod and myriocin). We compared this list of cell-wall/membrane-perturbation genes to all L2-mutant-linked differentially expressed genes, finding no significant overlap either globally (Fig. S4I), at the histone family level (Fig. S4J), or across individual L2 positions (Fig. S4K). Together, these results indicate that, independent of its role in transcription, the L2 has an additional regulatory capacity impacting actin and endocytosis, and the cellular periphery.

### Molecular origins of the L2’s regulatory complexity

Free, non-nucleosome-bound, histones are actively degraded by robust surveillance systems,^47,48^ making a cytosolic pool of L2 mutant histones unlikely to be key drivers of the non-chromatin GIs and phenotypes we observe. Thus, we wondered whether actin, endocytosis, and other membrane-associated GIs, despite their association with non-nuclear functions, might also be present in the nucleus. To identify such nuclear localization of GIs, we used four independent genome-wide datasets including GFP fusion protein localization^49^ and three independent chromatin fractionation studies,^50–52^ as well as sequence-based predictions like nuclear localization signals (NLSs)^53^ and annotated complexes^34^ (Fig. 4A; S5A). We implemented a weighted evidence score to integrate these diverse datasets and observed that, whereas the majority of L2 GIs across all mutants had either no or low supporting evidence, 61 L2 GIs had medium or high chromatin interaction confidence (Fig. 4B; Data S1). Because of their close association with chromatin, we name this class of GIs as anchor nodes, as they are most likely to be directly associated with chromatin. Anchor nodes were broadly similar to the overall distribution of GIs (Fig. 4C), with L2 positions 2 and 5 having the greatest number of anchor nodes. Further, anchor nodes were specifically enriched for acidic residues (*p*_adj_ < 0.001, Wilcoxon/Benjamini-Hochberg; Fig. 4D) and depleted for hydrophobic ones (*p*_adj_ < 0.01, Wilcoxon/Benjamini-Hochberg; Fig. 4E), consistent with their potential binding to lysine-rich histones.

**Figure 4.**
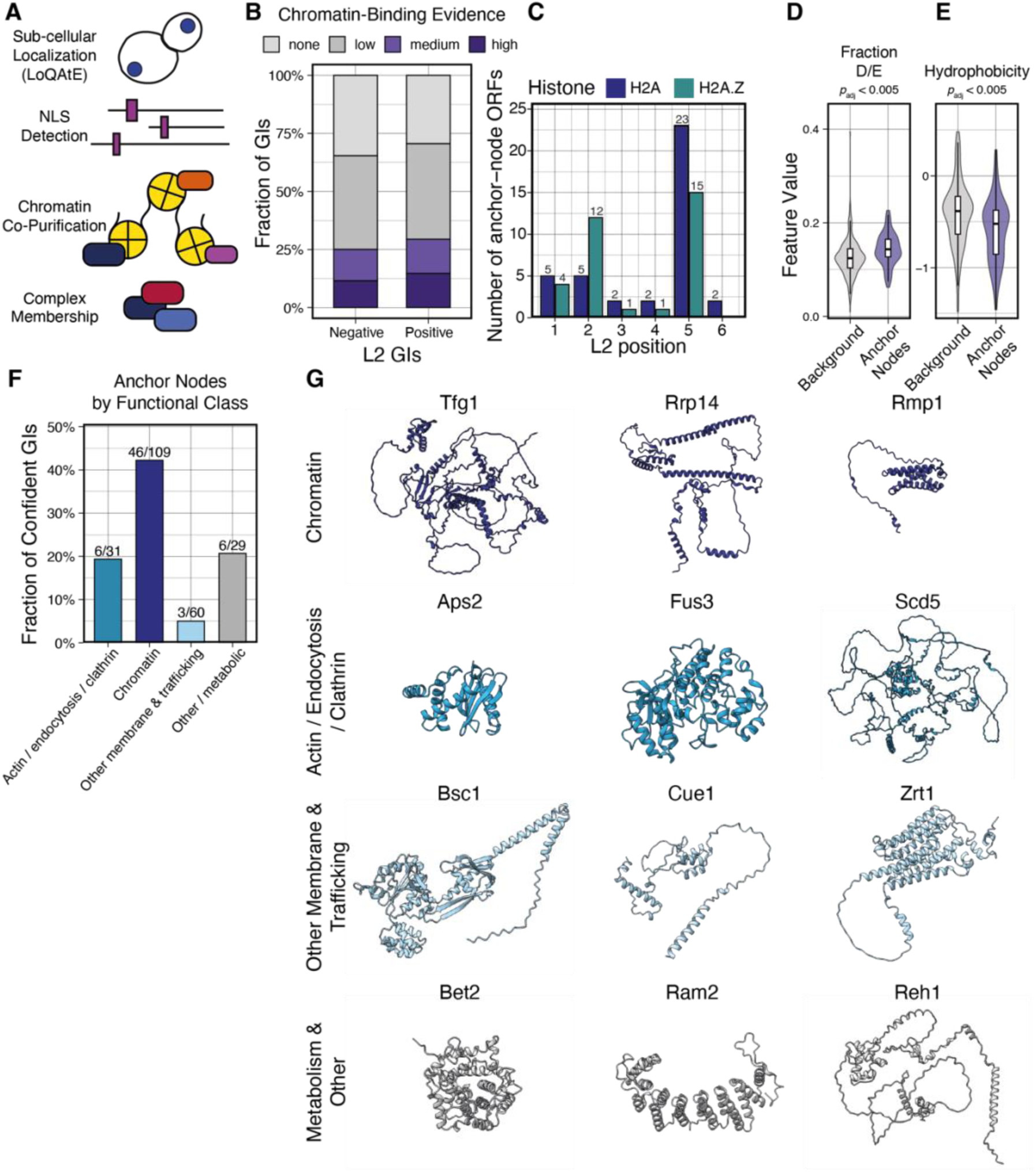
Molecular origins of the L2’s regulatory complexity. A, Scheme illustrating the input datasets for identifying chromatin-bound genetic interactors (GIs). Datasets include experimental sources including GFP-localization databases (LoQaTe^49^) as well as three independent chromatin fractionation studies, complex annotations (SGD), and NLS predictions. **B,** Distribution of anchor node confidence scores from low (no evidence) to high (3+ sources of evidence). **C,** Anchor node number by L2 position and histone. The number of anchor nodes is listed at the top of the corresponding bar. **D–E,** Biophysical properties of anchor nodes. (D), the fraction of negative amino acids (D/E) or, (E), the hydrophobicity (GRAVY). Nodes are compared to an equally sized pool of genes randomly picked from the rest of the genome. Statistical significance is assessed with a Wilcoxon rank test, and *p* was adjusted with the Benjamini-Hochberg procedure. **F,** Breakdown of anchor nodes across the four functional categories established in (4A–B) for all GIs in this study. **G,** AlphaFold3 predicted structures for example anchor nodes from (G) by functional class.

The majority of anchor nodes (75%) were directly related to chromatin, as expected (Fig. 4F). This included previously noted GIs related to transcription (e.g. Tfg1), but also nucleolus and ribosomal RNA processing (Rrp14, Rmp1) functions not previously associated with H2A and H2A.Z (Fig. 4G). Besides chromatin, we also identified several anchor nodes related to actin and endocytosis (6), membrane dynamics and trafficking (3) (Fig. 4F). Such non-chromatin anchor nodes were structurally and functionally diverse, including several intrinsically disordered proteins (Fig. 4G). Among actin and endocytosis nodes, they included clathrin-mediated endocytosis (Asp2), mating-related bud tip formation (Fus3), as well as cortical actin (Scd5). Membrane-related anchor nodes included cell-surface protein Bsc1, ER-membrane bound ubiquitin ligase Cue1, as well as zinc transporter Zrt1. Finally, anchor nodes also included geranylgeranyltransferases, Bet2 and Ram2, as well as the cytosolic ribosome biogenesis factor Reh1. Together, these results suggest that, whereas the L2’s function is closely connected to chromatin and transcription, the origins of its phenotypic impact also encompass molecular associations with cytosolic functions.

## DISCUSSION

Using the H2A/H2A.Z L2 as a model, we demonstrate that the nucleosome core encodes substantial phenotypic variation—including a large proportion of beneficial states. Such phenotypic variation is linked to corresponding regulatory complexity, and single L2 point mutations have distinct, robust regulatory interactions. Further, these L2 regulatory interactions are not confined solely to classical chromatin regulatory factors such as transcription, but include diverse processes from actin and endocytosis to membrane trafficking. Together, our results link the nucleosome core to potentially beneficial functional variation and highlight a deeper connection between the nucleosome and functions of the cellular periphery than previously appreciated.

Although the evolution of H2A variants is still debated,^1,2,13,15^ our findings posit that, in addition to structural constraints, network-level interactions could have also been a key driving force shaping H2A evolution. Whereas the phenotypic landscape and interaction network of H2A is large, that of H2A.Z is narrower and more specifically constrained by a discrete biochemical space. This would be consistent with H2A.Z specializing from a more generalist H2A ancestor, acquiring key nodes within the larger network that could have propelled its divergence from the ancestral H2A before reaching the evolutionary stasis^2,13^ observed in extant eukaryotic genomes. This would predict that H2A’s additional lineage-specific variants,^54–59^ including macroH2A and H2A.W, likely subdivided the ancestral H2A network into further specialized roles. Future regulatory maps of H2A, H2A.Z, and lineage-specific histone variants in diverse species will reveal whether this is the case, and if so which regulatory functions––whether chromatin and transcription, or broader functions at the cellular periphery––have been crucial for shaping their evolution.

It has been proposed that the ancestral function of histones is to package, protect and regulate access to the genome, for example in the context of transcription.^60^ Whereas our results echo these roles for the nucleosome in cellular physiology, they reveal that the nucleosome’s impact also includes links to functions at the cellular periphery. Such a relationship could have evolved before the nuclear compartment, when the genome was already highly chromatinized with histones.^61^ This would predict that many of the shuttling mechanisms that we identify linking non-chromatin proteins to the nucleosome could reflect an evolved mechanisms to preserve functional links despite the creation of a membrane barrier. Consistent with this interpretation, many of the anchor nodes we identified––particularly relating to actin and endocytosis––appear to have arisen in the last eukaryotic common ancestor. However, given that the boundary between Eukarya and Archaea is rapidly narrowing with the discovery of Asgard lineages,^62,63^ we anticipate that future phylogenomic insights, combined with cross-species functional comparisons, will clarify the interplay between the evolution of nucleosomes, their regulatory networks, and the nucleus during eukaryogenesis.

Finally, the recruitment of diverse cellular functions at the nucleosome surface prompts the question as to how coupling divergent processes contributes to phenotypic innovation and evolutionary trajectories. Complex traits often arise through the convergence of multiple genetic features. This has been demonstrated by theory,^64,65^ by observing natural populations,^66–68^ and by experiment.^69,70^ Although regulatory changes to chromatin can give rise to new and phenotypically beneficial variants,^71^ whether such changes are a driver of evolvability, and if so what their molecular basis remains an open question. The density of the nucleosome’s regulatory network that we uncover creates potential for complex traits to converge to drive rapid evolution; only a single amino acid change is required to rewire functions across the cell, creating or destroying molecular links with consequence from gene expression to endocytosis. Addressing these questions will not only transform our understanding of how, despite being deeply conserved, core eukaryotic machinery has shaped evolution, but also how these mechanisms can be coopted by disease to fuel pathology.

## ACKNOWLEDGEMENTS

We thank the entire Berger group, as well as Bassam Al-Sady, for their insightful, considerate, and helpful discussions. We especially thank Carla Brillada and the Ramundo group, as well as Zdravko Lorković, Elin Axelsson-Ekker, and Chung Hyun Cho for their invaluable assistance with high-throughput robotic equipment, preparing antibodies, biochemical assays, and bioinformatic analyses. We further thank the Molecular Biology Service and Media Kitchen for a constant supply of plates, basic reagents, cloning and sequencing services, and general technical assistance. Additionally, we thank the Vienna BioCenter Core Facilities, in particular the Next Generation Sequencing facility for their advice and swift handling of all our requests. This work was supported by core funding from the Gregor Mendel Institute, Austrian Academy of Sciences (F.B.); grants from die Österreichischer Wissenschaftsfonds FWF (TAI304 and ESP213, F.B. and Z.H.H., respectively); and an EMBO Postdoctoral Fellowship (ALTF169-2020, Z.H.H.). For open access purposes, the authors have applied a CC BY public copyright license to any author-accepted manuscript version arising from this submission.

## METHODS

### Study Organisms and General Culture Procedures

Yeast were cultured under standard conditions, 30 °C, for *Saccharomyces cerevisiae* (BY4741, BY4742). Unless otherwise stated, growth media used were either CSM (Complete Synthetic Media) or YPD (Yeast Extract, Peptone, Dextrose).^73^

### Strain Manipulation

*S. cerevisiae* strains were manipulated either using CRISPR^74^ with HR templates synthesized by commercial vendors (IDT), or through classical PCR-directed homologous recombination.^73,75^ Heat-shock was used to transform chemically-competent parent strains.^76^ Donor templates for PCR-directed homologous recombination were prepared using either Gibson assembly^77^ using reagents supplied by the IMP Molecular Biology Service combined, with site directed mutagenesis, or by direct synthesis from commercial vendors (IDT). All manipulations were verified by PCR genotyping, and where appropriate sequencing of the relevant locus.

### Phenotypic Assays

Phenotypic assays in this study were performed using a plate-based bulk growth assay. OD_600_ was monitored in 384-well plates (Nunc) continuously under the stress conditions indicated in figure legends at the standard growth temperature with shaking. For all yeast assays, saturated cultures were sub-cultured 100-fold and all assays were performed in organism-appropriate growth medium. Data collection was performed on either a BioTek Epoch2 with a Biostacker or Synergy4 instrument. Area under the curve (AUC) was calculated from these growth curves in R (v4.1.3) using the Rstudio IDE (2022.12.0+353) using the Growthcurver package v0.3.1 (https://cran.r-project.org/web/packages/growthcurver/vignettes/Growthcurver-vignette.html). All subsequent data analysis was done in R.

### Transcriptomic Profiling

Total RNA was isolated using the IMP Molecular Biology Service RNA bead isolation kit, following the included protocol with a modified tissue-lysis procedure. Cell pellets from 5 ml mid-long cultures (OD600 ∼ 0.5) were lysed at room temperature in the supplied guanidine thiocyanate/Triton X-100 lysis buffer, and mechanically disrupted using glass beads in a Retsch MM400 mixer mill. Following clarification of lysates, RNA was purified using paramagnetic beads and a KingFisher Flex 96 automated pipeline. Total RNA was quantified by Nanodrop, and quality control was performed by agarose gel electrophoresis.

3‘Tag-seq libraries were prepared in 96-well plates as described previously.^78^ Total RNA (80 ng in 4 µl) was combined with 1 µl of a 0.2 µM barcoded oligo-dT reverse-transcription primer containing a 7-bp UMI, heated at 72°C for 3 min, and immediately cooled. Reverse transcription was performed using SmartScribe reverse transcriptase (Takeda). Barcoded cDNA samples were pooled and purified using a 1.2× SPRI cleanup. The purified DNA–RNA substrate was tagmented using pre-loaded Tn5 transposase at 55°C for 8 min, and the reaction was terminated with SDS. Following a 2× SPRI cleanup, libraries were amplified with KAPA HotStart ReadyMix and P5/P7 primers for 12 PCR cycles. Final libraries were purified using a 0.8× SPRI cleanup, eluted in 10 mM Tris-HCl (pH 8.0), and stored until sequencing on either an Illumina NovaSeqX 10B. Data were demultiplexed and processed for analysis using a Nextflow pipeline adapted from Bourguet et al. 2025,^79^ using the *S. cerevisiae* R64-1-1 reference genome. Differential expression analysis was performed in R using DESeq2.^72^

### Quantitative Genetic Interaction Mapping

Genetic interactions for all 21 H2A and H2A.Z L2 mutants was performed essentially as described previously.^35^ Briefly, each histone mutation was introduced to a MATalpha query strain (YZH_1419/Y#7092^35^; S288C can1delta::STE2pr-Sp_his5 lyp1delta his3delta1 leu2delta0 ura3delta0 met15delta0) along with a natMX cassette for selection. These were then crossed to MATa libraries of either the yeast gene knockout collection^37^ (Horizon Discovery) or haploid Decreased Abundance by mRNA Perturbation^38^ (DAmP; Horizon Discovery) using robotically assisted (Singer Instruments ROTOR+) pinning in 1,536-spot arrays in biological duplicate. Diploids were selected by passaging on growth medium supplemented both with G418 (200 mg/L; Enzo) and nourseothricin (100 mg/L; Jena Biosciences) before sporulation on rich SPO^35^ medium for 5 days. Double mutants were then isolated by sequential passaging on growth medium to select for MATa-specific marker expression (CSM–His), and then selection for both MATa marker and both antibiotic selection cassettes for the histone (natMX) and gene disruption (kanMX). Double mutants were initially pinned onto YPD and grown for 16 hrs before initial colony size quantification (Singer Instruments PhenoBooth). These were then pinned onto three different conditions in parallel: YPD, or supplemented with 0.5 M NaCL, 50 µg/ml Doxorubicin (Med Chem Express). YPD plates without any additives were transferred to an incubator set to 39 °C, and all plates were grown for 16 hrs before final colony area quantification.

Colony sizes from the SGA screen were used as the quantitative fitness readout. After removing empty and unscored colonies (missing size or Size < 10 px), colony size was log-transformed (Size_log) and used as the Gaussian response. Each colony was annotated with plate-position covariates (row, column, edge status, pinning order and a pin-missing flag) so that spatial and pinning artefacts could be regressed out.

Genetic interactions (GIs) were estimated with a single joint generalized linear mixed model fitted in glmmTMB (Gaussian family, maximum likelihood, REML = FALSE). For colony *c* carrying gene deletion (ORF) *o*, histone backbone *h*, substitution *s* and environment *e*, the log colony size *Y_c_* was modelled as:

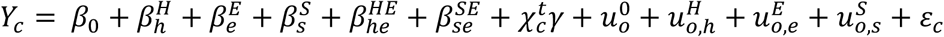

with

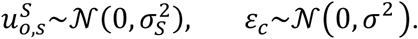

Where *β* terms are for fixed effects encompassing histone and environmental variables,*χ^t^_c_γ* collects the plate/pinning nuisance covariates, and *u* terms capture per-ORF random effects as they depend on histone, environment, or mutation. The genetic interaction for substitution *s* in the presence of query gene *o* is the corresponding conditional mode (BLUP) of the per-ORF substitution random effect:

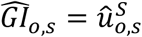

i.e. the ORF-specific departure of the double-perturbation phenotype from the additive expectation defined by the population and main effects. From these GI’s, contrasts for a given L2 mutant relative to the respective WT control strains were calculated as:

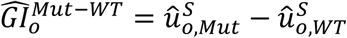

From these GIs, A Wald z-statistic (estimate/SE) and two-sided was computed per interaction, and used to set a threshold for significant interactions. Unless otherwise noted, |GI| > 0.3 and a |z-statistic| > 2.5 was used as a stringent cutoff. These values were empirically determined after testing a range of cutoffs, generating STRING^43^ (confidence > 400) networks for each and selecting the settings with the highest network significance and biological coherence.

### Anchor Nodes Identification

To classify genetic-interaction partners as candidate chromatin-anchored nodes we implemented an additive evidence score built from four independent annotation sources. For every ORF in the genome we defined four binary indicators: (i) *e*_loc_, nuclear or nucleus-enriched GFP localisation in LoQAtE^49^; (ii) *e*_chrom_, detection in at least one chromatin-proteomics dataset (union of the compiled datasets, ≥1 dataset)^50–52^; (iii) *e*_cplx_, membership in a protein complex annotated as nuclear^34^; and (iv) *e*_nls_, a NLStradamus^53^ posterior ≥ 0.8. ORFs absent from a given resource were scored 0 for that component, and coverage was recorded separately so that missingness is distinguishable from negative evidence in the supplementary table.

Because the four sources differ greatly in prior frequency, components were weighted by their information content (surprisal). For source *j* with genome-wide marginal frequency *p_j_*, the weight was *w_j_* = −log_2_ (*p_j_*) bits, and the score for ORF *i* was *E_i_* = Σ*_j_ w_j_*·*e_ij_*. Marginals were estimated on the full scored frame (*n* = [N] ORFs); the resulting weights were *w*_loc_, *w*_chrom_, *w*_cplx_, and *w*_nls_ bits, so that rare evidence (nuclear complex membership, *p*_cplx_) contributes several-fold more than common evidence (chromatin proteomics, *p*_chrom_). ORFs were additionally binned into evidence tiers (none, low = 1 source, medium = 2, high = ≥3), and an ORF was called a chromatin-anchored node if it was supported by ≥ 2 independent sources.

### Quantification and Statistical Analysis

All statistical tests were performed in R, with all details regarding specific tests applied included in the appropriate figure legend. Unless otherwise noted, all enrichment calculations were done with a Fisher’s exact test, and corrected following the Benjamini-Hochberg procedure where appropriate. All tests of differences of mean were Wilcoxon rank sum tests, also corrected by Benjamini-Hochberg where appropriate and unless otherwise noted.

### Material Availability and Contact Information

All strains, data, analysis scripts, and other materials and computational resources generated by this study are available upon request to the lead contact. Transfer of strains and other biological materials generated by this study require the completion of a materials transfer agreement.

---

**Figure S1.**
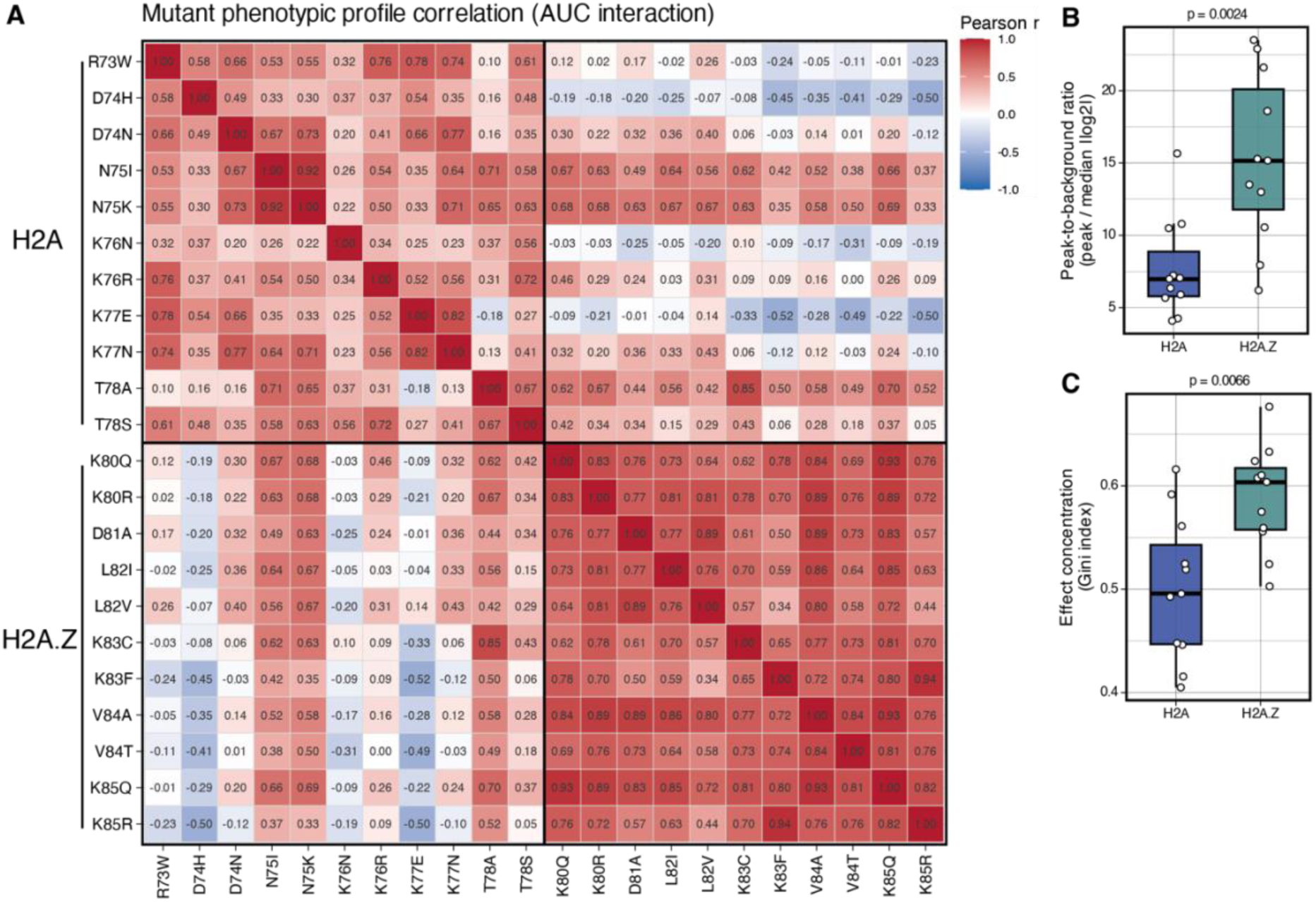
Related to figure 1. **A,** Correlation matrix for phenotypes from (1D). **B–C,** Alternative specialization indices for H2A.Z vs. H2A demonstrating robustness. (B) Comparison of the highest growth advantage relative to the median phenotypic severity. (C) Gini index for all phenotypes.

**Figure S2.**
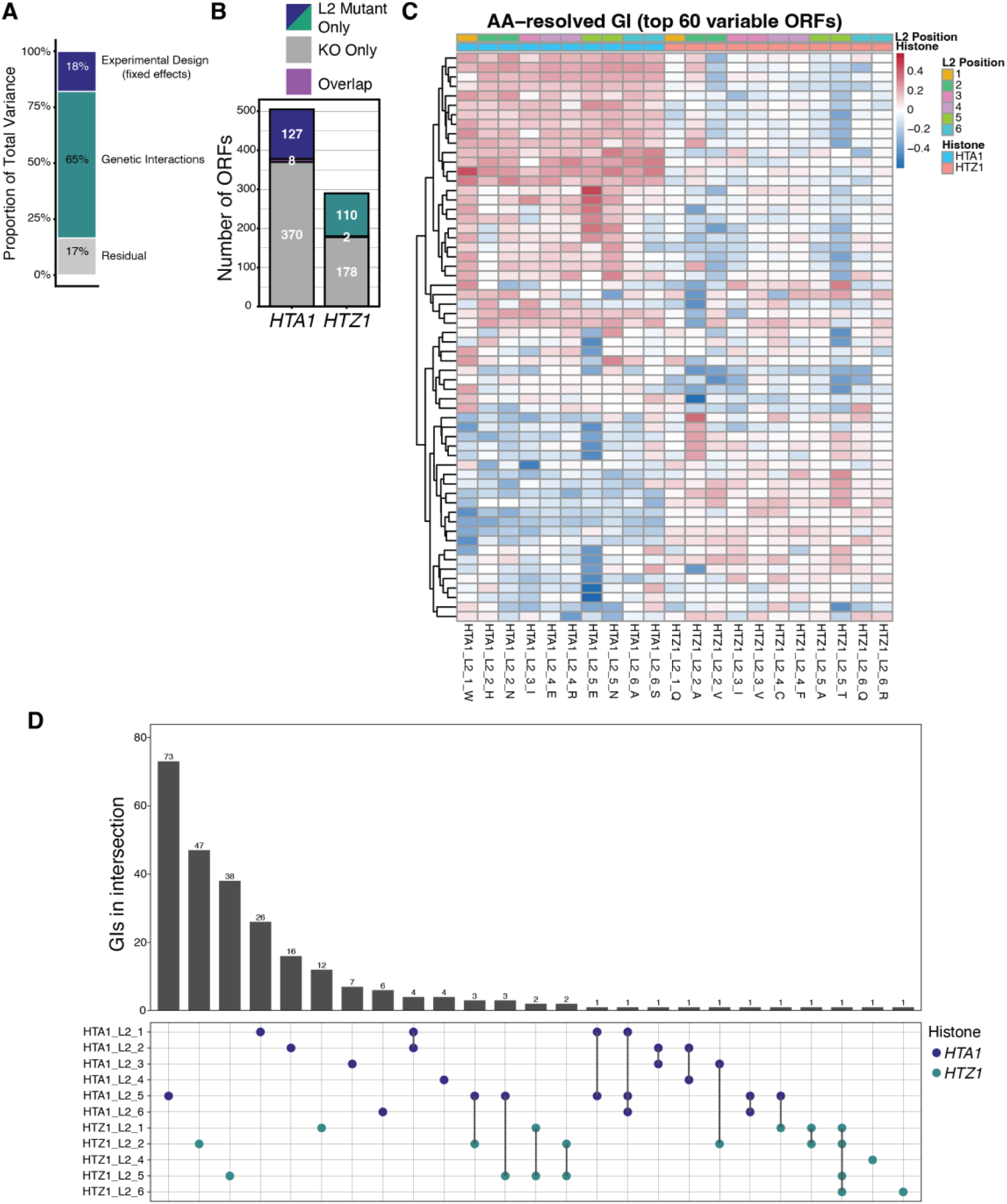
Related to figure 2. **A,** Breakdown of the contribution of each linear mixed effect model to measured genetic interactions (GI). **B,** Comparison of GIs identified in this study to those from The Cell Map for *hta1*Δ (H2A) and *htz1*Δ (H2A.Z).^40^ Genetic interactions are all negative, and are defined as in Costanzo et al.^39^ (ε <-0.08 and *p* < 0.05). **C,** Top 60 most variable GIs across all L2 mutants. **D,** Upset plot describing the overlap between GIs across all L2 positions in our study.

**Figure S3.**
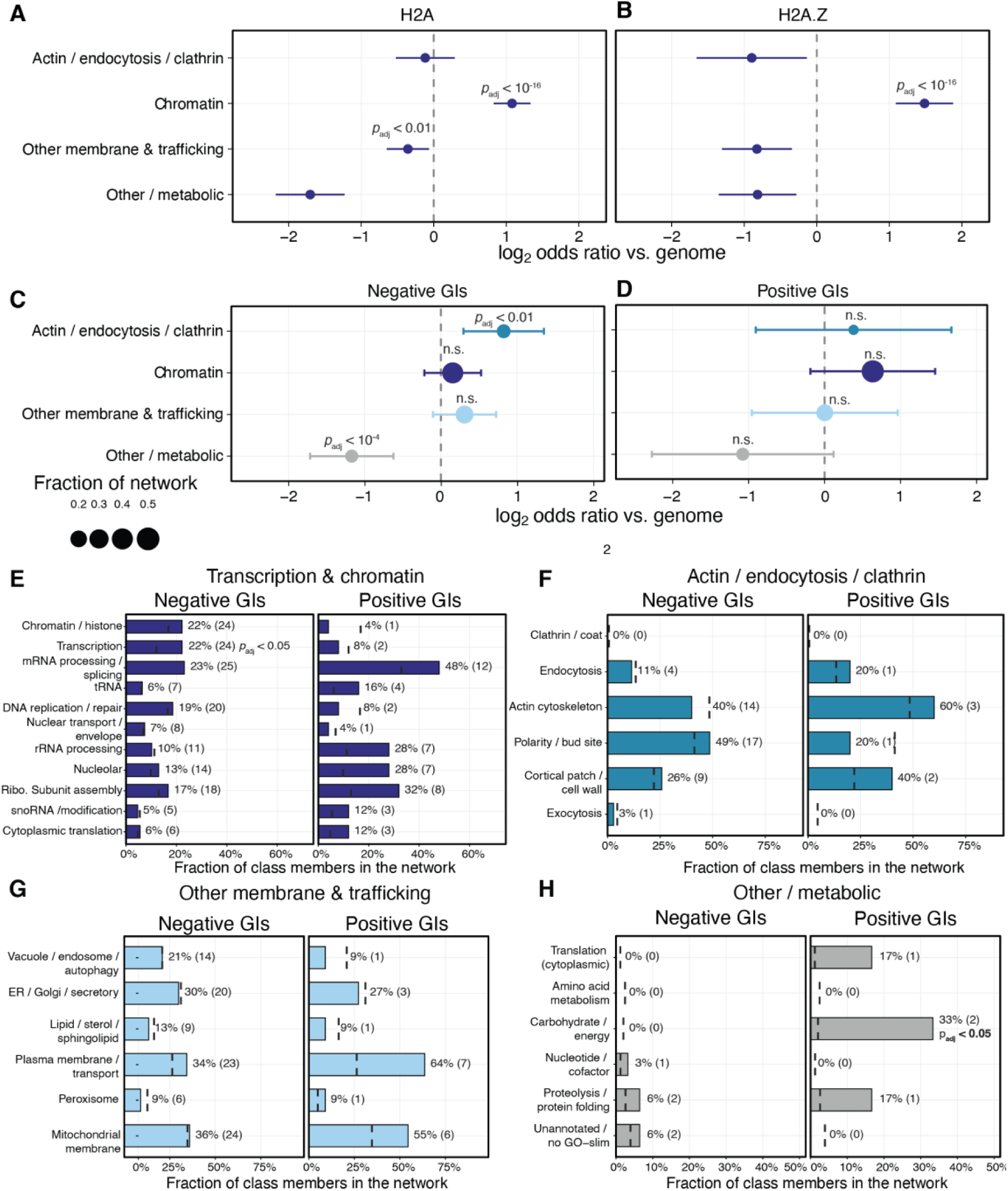
Related to figure 3. **A–B,** Enrichment analysis across four key functional categories for genetic interactions for *hta1*Δ and *htz1*Δ from The Cell Map.^40^ Genetic interactions are all negative, and are defined as in Costanzo et al.^39^ (ε <-0.08 and *p* < 0.05). **C–D,** As in (A–B), but for negative (C) and positive (D) GIs from this study. **E–H,** As in (A–B), but decomposed into sub-categories for transcription and chromatin (E), actin and endocytosis (F), other membrane-related proteins (G), and other or metabolic proteins (H). Unless otherwise noted, statistical significance is assessed with a Fisher’s exact test with a Benjamini-Hochberg adjustment.

**Figure S4.**
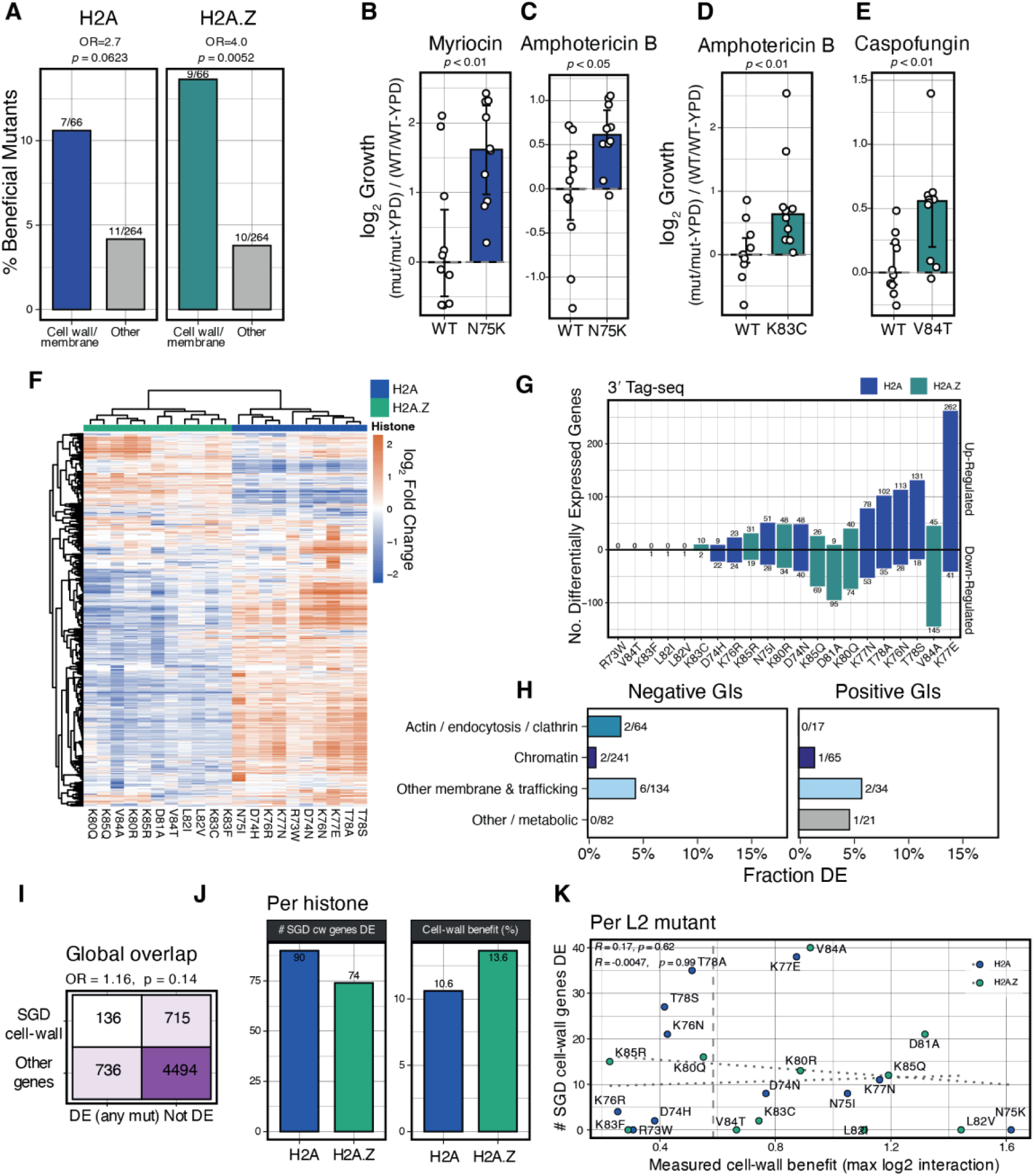
Related to figure 3. **A,** Enrichment of phenotypically beneficial L2 growth phenotypes by category. Cell wall/membrane conditions are defined as: amphotericin B, caspofungin, clotrimazole, fingolimod, and myriocin. Phenotypically beneficial mutants are defined as having a growth advantage of at least 50% relative to WT. Statistical significance is assessed with a Fisher’s exact test. **B–E,** Example growth phenotypes for the indicated H2A (B–C) and H2A.Z (D–E) L2 mutants in selected cell-wall/membrane-related stress conditions. Growth is normalized to the median of the WT. Statistical significance is assessed with a Wilcoxon rank test. **F,** Hierarchically clustered heatmap of 3’ Tag-seq results for all 21 L2 mutants across both H2A and H2A.Z for all genes. Scale is the log2 Fold Change calculated with DESeq2.^72^ **G,** Number of differentially expressed genes per L2 mutant. Differential expression is defined as |log_2_ Fold Change| > 1 and *p*_adj_ < 0.05. **H,** Breakdown of the number of total negative or positive GIs that are differentially expressed, matched by L2 mutant. **I–K,** Comparison of L2-mutant differentially expressed (DE) genes with those annotated as having a phenotype in any of the cell wall/membrane-related stress conditions (amphotericin B, caspofungin, clotrimazole, fingolimod, and myriocin). Annotations were taken from phenotypic data stored in the Saccharomyces genome database (SGD).^34^ Comparisons are done either at the level of all GIs (I), broken down by histone (J), and for individual L2 mutants (K).

**Figure S5.**
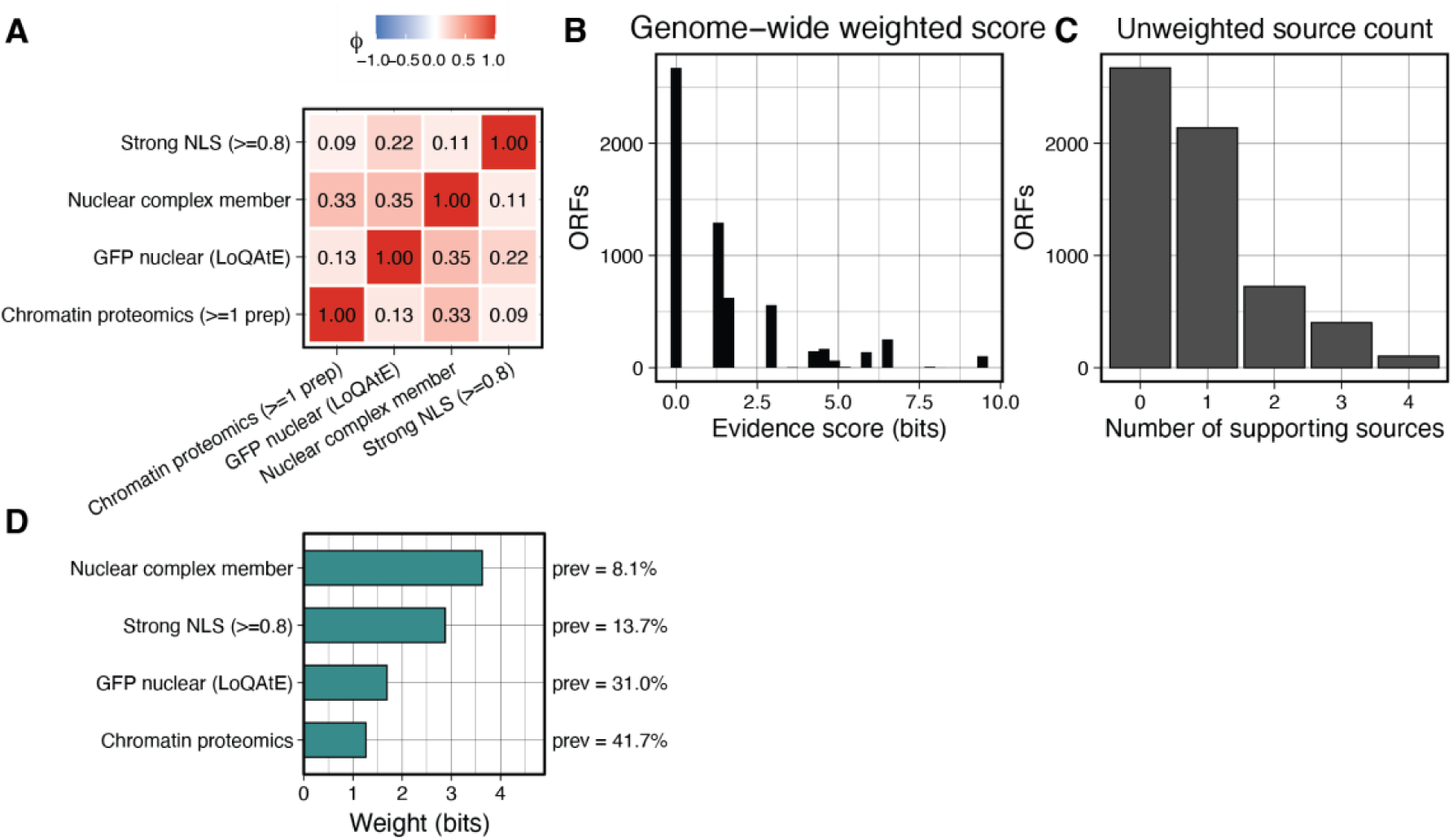
Related to figure 4. **A,** Genome-wide prevalence of the four data sources used to build anchor node confidence scores. **B,** Mean-square contingency coefficient for the four input data sources used to assess anchor node confidence. **C,** Genome-wide confidence of chromatin binding from all four integrated data sources. Evidence score is the sum of weighted evidence for a given ORF, and is displayed as a histogram of all 6039 annotated ORFs in the *S. cerevisiae* genome. **D,** Histogram of the number of data sources supporting each anchor node call.

